# Bacterial succession, functional organization, and ecological drivers during spontaneous ukwa (*Treculia africana*) fermentation

**DOI:** 10.64898/2025.12.29.696824

**Authors:** Kelechi Stanley Dike, Chidinma Francisca Ezeh, Monicah Maseke, Chinwe Happiness Justice-Alucho, Chibundu N Ezekiel

**Affiliations:** Department of Microbiology, Imo State University, Owerri, Nigeria; Department of Food Science and Technology, Federal University Owerri Nigeria; Department of Health Sciences, Machakos University Kenya; Biological Sciences, Clemson University USA; BOKU University, Institute for Bioanalytics and Agro-Metabolomics, Department of Agricultural Sciences, Tullin, Austria

**Keywords:** Ukwa, *Treculia africana*, Bacterial succession, 16S rRNA gene sequencing, Bacterial ecology, Solid-state fermentation, Bacterial community dynamics

## Abstract

Spontaneous fermentation of ukwa (*Treculia africana*) involves complex microbial succession that remains poorly characterized. Here, bacterial community dynamics during ukwa fermentation were resolved across six time points using high-throughput 16S rRNA gene sequencing. Fermentation followed a structured, stage-wise trajectory, beginning with plant-associated aerobic taxa, transitioning through a mid-phase dominated by fermentative firmicutes, and culminating in late-stage enrichment of acid-tolerant lactic and acetic acid bacteria. Microbial richness increased progressively during fermentation, while community composition clustered by time and correlated strongly with declining pH and moisture. Multivariate analyses identified these physicochemical gradients as primary drivers of succession. Network analysis further revealed distinct co-occurrence modules, indicating coordinated microbial guilds that underpin functional transitions during fermentation. This study provides the first sequencing-based ecological framework for ukwa fermentation and highlights how environmental filtering and microbial interactions shape community assembly in traditional African solid-state fermentations. These insights provide a basis for improving process control and guiding future starter culture development for ukwa and related fermented foods.

## 1. INTRODUCTION

Ukwa (*Treculia africana*) is a traditionally fermented seed product widely consumed in southern Nigeria, contributing to dietary diversity and local food security (Hill et al., 2025; Ojimelukwe & Ugwuona, 2021). The seeds are nutrient-dense, providing substantial protein, minerals, and bioactive compounds, underscoring their nutritional importance in local diets (Oyetayo & Oyetayo, 2020). Spontaneous solid-state fermentation enhances sensory qualities, improves nutrient accessibility, and reduces antinutritional factors in plant substrates (Pandey et al., 2000). Despite these advantages, product quality often fluctuates, as the microbial ecology underlying these indigenous fermentations remains only partially understood (Azokpota et al., 2006; Parkouda et al., 2009). Previous culture-based studies of African seed fermentations consistently report Bacillus species, lactic acid bacteria (LAB), and acetic acid bacteria (AAB) as major contributors to acidification, flavour development, and proteolysis within these systems (Ouoba et al., 2003; Parkouda et al., 2009; Achi, 1992). More broadly, LAB have long been recognized as central starter and adjunct cultures across various food fermentations, playing crucial roles in acidification, inhibition of spoilage organisms, and the generation of key flavour and textural attributes (Leroy & De Vuyst, 2004). Preliminary culture-based work on ukwa by the present author identified *Lactobacillus* and *Bacillus* among the dominant genera (Dike, 2024), but such approaches capture only a fraction of the microbial community, offering limited ecological insight (Ercolini, 2017). High-throughput 16S rRNA gene sequencing enables comprehensive characterization of microbial succession and its environmental drivers, as shown for toddy fermentation and other traditional plant-based foods (Das & Tamang, 2021; Tamang et al., 2020). Solid-state fermentations often exhibit deterministic microbial structuring driven by oxygen depletion, progressive acidification, and declining moisture, resulting in predictable microbial transitions (Hao et al., 2021; Yan et al., 2025).

Mapping these ecological dynamics is crucial for enhancing process control, improving product consistency, and guiding the development of starter cultures tailored to local substrates. Despite the cultural and nutritional relevance of ukwa, its microbial ecology remains uncharacterized. No sequencing-based studies have examined its bacterial community structure, successional patterns, or environmental determinants. This study presents the first high-throughput analysis of bacterial succession during ukwa fermentation using 16S rRNA gene sequencing. By integrating microbial profiling with pH and moisture measurements, we identify core taxa, key ecological transitions, and stage-specific functional modules. Taken together, these findings establish a microbiological baseline for ukwa fermentation and provide mechanistic insights that are relevant for improving fermentation reliability and guiding the development of tailored starter cultures.

## 2. MATERIALS AND METHODS

### 2.1 Sample Collection and Physicochemical Analysis

Approximately eight mature green fruits of *Treculia africana* (totalling ∼40 kg) were collected in July 2024 from Orodo, Imo State, Southeast Nigeria (5°36′59.7″ N, 7°02′11.0″ E). A single batch of solid-state fermentation (SSF) was initiated by heaping the fruits at ambient temperature (28–30°C) for eight days. Fermenting samples (∼40 g) were collected at 0, 24, 48, 96, 144, and 192 h by pooling equal portions from multiple surface and interior positions within the fermenting heap to generate one biologically representative composite sample per time point. This ecological time-series of a single traditionally fermented ukwa batch establishes a baseline microbiome profile using high temporal resolution, an approach widely applied in first-generation microbiome studies of under-characterized fermented foods. Samples were sequenced alongside cassava fermentation samples as part of a broader study of Nigerian fermented foods (BioProject PRJNA681567); cassava fermentation results were reported separately (Dike et al., 2022). For physicochemical analysis, pH was measured using a 1:10 (w/v) seed slurry prepared in distilled water and recorded with a calibrated Hanna HI98103 pH meter. Moisture content was determined by oven-drying samples at 105°C to a constant weight, following the AOAC Method 925.10. All measurements were performed in triplicate. Temporal changes in pH and moisture were evaluated using simple linear regression in R (v4.5.1).

### 2.2 DNA Extraction and Quantification

Genomic DNA was extracted using the ZymoBIOMICS™ DNA Miniprep Kit (Zymo Research, Irvine, CA, USA). Approximately 10 g of each fermentation sample was mixed with 5 mL of chilled ultrapure water and vortexed thoroughly. The mixture was centrifuged at 800 × g for 1 min to pellet solids, and the supernatant was collected. This washing step was repeated twice. The pooled supernatant (∼15 mL) was centrifuged at 12,000 × g for 5 min to pellet free microbial cells. DNA was extracted from the resulting pellet according to the manufacturer’s instructions. DNA concentration was determined using a Qubit™ 3.0 Fluorometer (Invitrogen, USA), and integrity was assessed by 1% (w/v) agarose gel electrophoresis.

### 2.3 Amplicon Library Preparation and Sequencing

Amplicon libraries targeting the bacterial V3–V4 region (∼450 bp) of the 16S rRNA gene were prepared using primers 357F (5′-CCTACGGGNGGCWGCAG-3′) and 806R (5′-GACTACHVGGGTATCTAATCC-3′) as described by Klindworth et al. (2013). Primers included Illumina overhang adapter sequences. PCR amplicons were indexed using Nextera XT adapters and purified with AMPure XP magnetic beads (Beckman Coulter, USA). Final libraries were pooled with a 10% PhiX control and sequenced on an Illumina MiSeq platform (2 × 300 bp paired-end reads) at Laragen Inc. (Culver City, CA, USA).

### 2.4 Bioinformatics and Data Processing

Raw paired-end Illumina MiSeq reads were processed by Laragen Inc. (Culver City, CA, USA) using QIIME 2 (version 2024.2; Bolyen et al., 2019). Quality assessment was performed using FastQC (Andrews, 2010), followed by quality filtering, denoising, paired-end merging, and chimera removal using the DADA2 plugin (Callahan et al., 2016). Amplicon sequence variants (ASVs) were generated using the DADA2 denoising algorithm, and singleton ASVs were removed to reduce sequencing noise and artifacts. Taxonomic classification was performed using the QIIME 2 feature classifier trained on the SILVA v132 reference database (Quast et al., 2013). Sequences assigned to Archaea, mitochondria, or chloroplasts were removed prior to downstream analysis. The final ASV feature table containing 431 ASVs and corresponding taxonomic assignments was exported from QIIME 2 and imported into R (v4.5.1) using the phyloseq package (McMurdie & Holmes, 2013) for ecological and statistical analyses.

### 2.5 Ecological and Statistical Analysis

All analyses were performed in R (v4.5.1) using the phyloseq package (McMurdie & Holmes, 2013). Descriptive statistics were used to summarize read counts and ASV richness. To account for uneven sequencing depth across samples (range: ∼37,000–230,000 reads per sample), ASV count data were rarefied to the minimum library size prior to calculation of alpha-diversity metrics (observed ASV richness, Shannon diversity index, Simpson’s diversity index), following established protocols for normalizing microbiome count data (McMurdie & Holmes, 2014; Schloss, 2024). Rarefaction curves were generated to confirm sufficient sequencing coverage for capturing bacterial community diversity (Supplementary Fig. S2). Taxonomic composition analyses (relative abundances at phylum and genus levels; Figs. 2–3) were performed on unrarefied count data, as relative abundance transformations inherently account for sequencing depth differences (Gloor et al., 2017). Alpha-diversity indices were compared across fermentation time points. Beta diversity was assessed using Principal Component Analysis (PCA) on genus-level relative abundance data. Spearman correlations were computed to examine relationships between environmental variables (pH, moisture) and microbial metrics. Differentially abundant taxa across fermentation stages were identified using LEfSe (Segata et al., 2011). Microbial co-occurrence networks were constructed with the NetCoMi package (Peschel et al., 2021) using a Spearman correlation threshold of 0.01. Network modularity was calculated using the cluster_fast_greedy algorithm in igraph (Csárdi & Nepusz, 2006) and visualized in Gephi (version 0.10.1; Bastian et al., 2009). Functional predictions were generated using FAPROTAX (Louca et al., 2016).

**Fig. 1.**
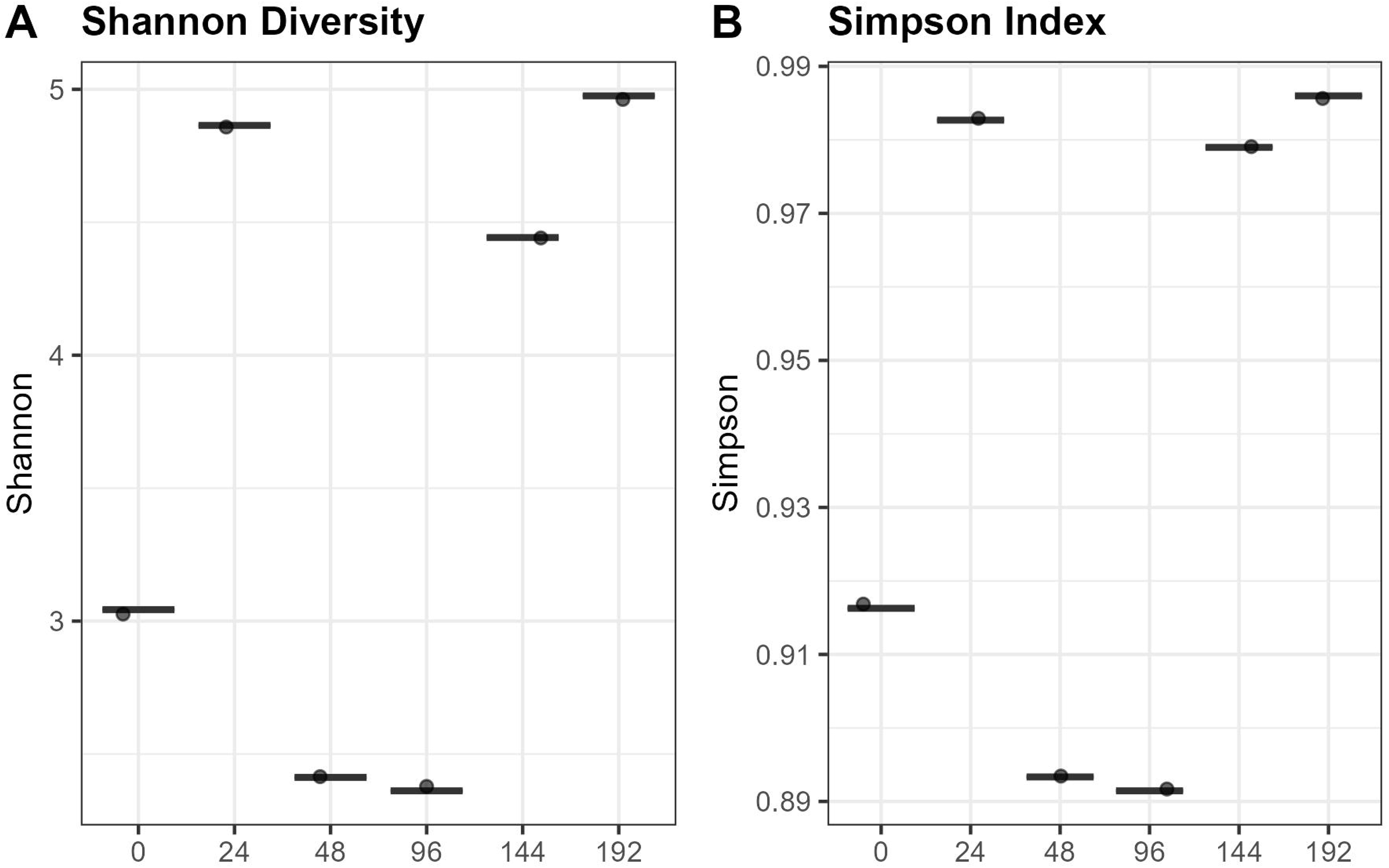
Alpha-diversity indices during ukwa fermentation. (A) Shannon diversity and (B) Simpson evenness across six time points (0–192 h).

**Fig. 2.**
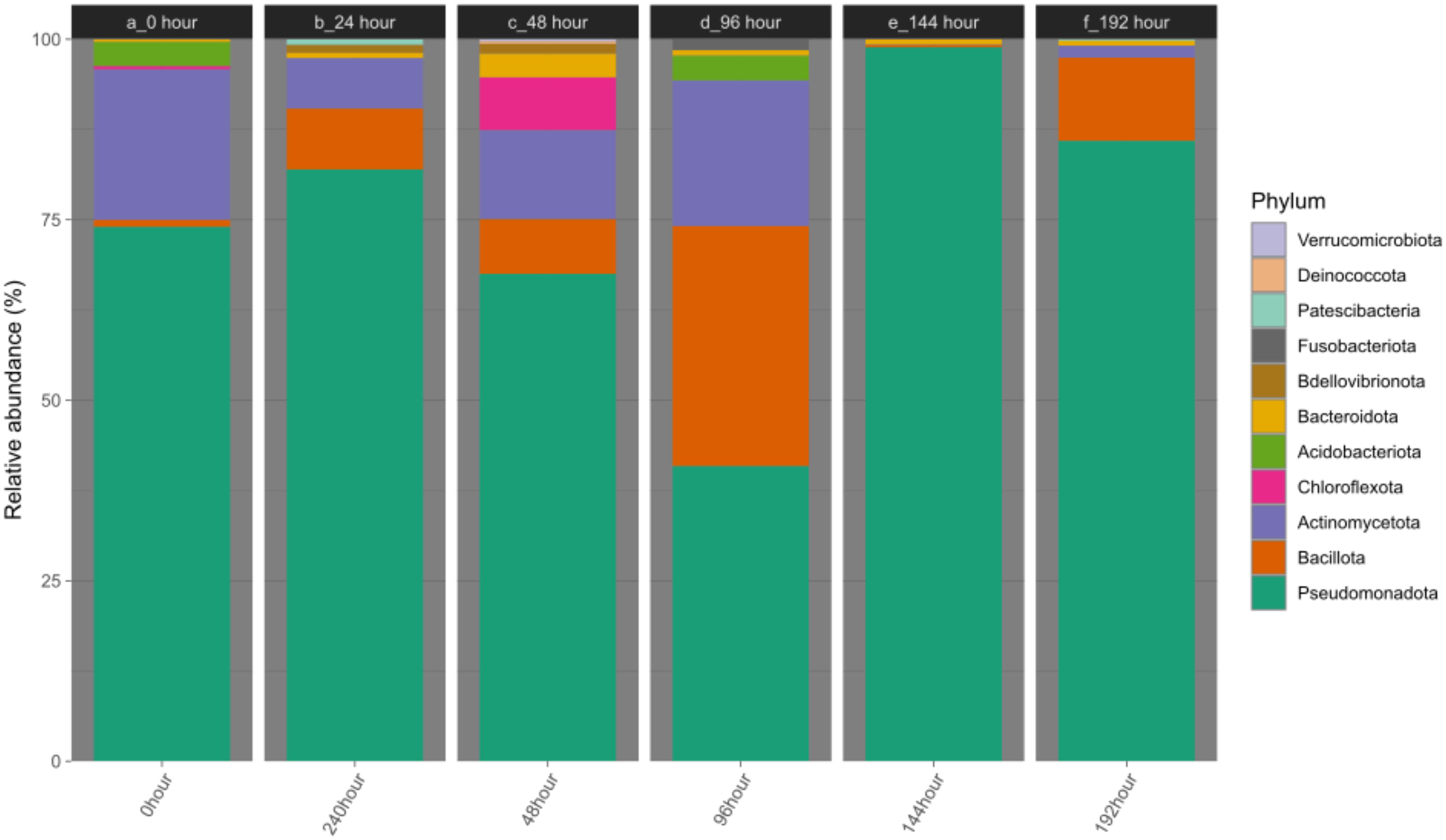
Phylum-level bacterial composition during ukwa fermentation. Relative abundances across six time points (0–192 h).

**Fig. 3.**
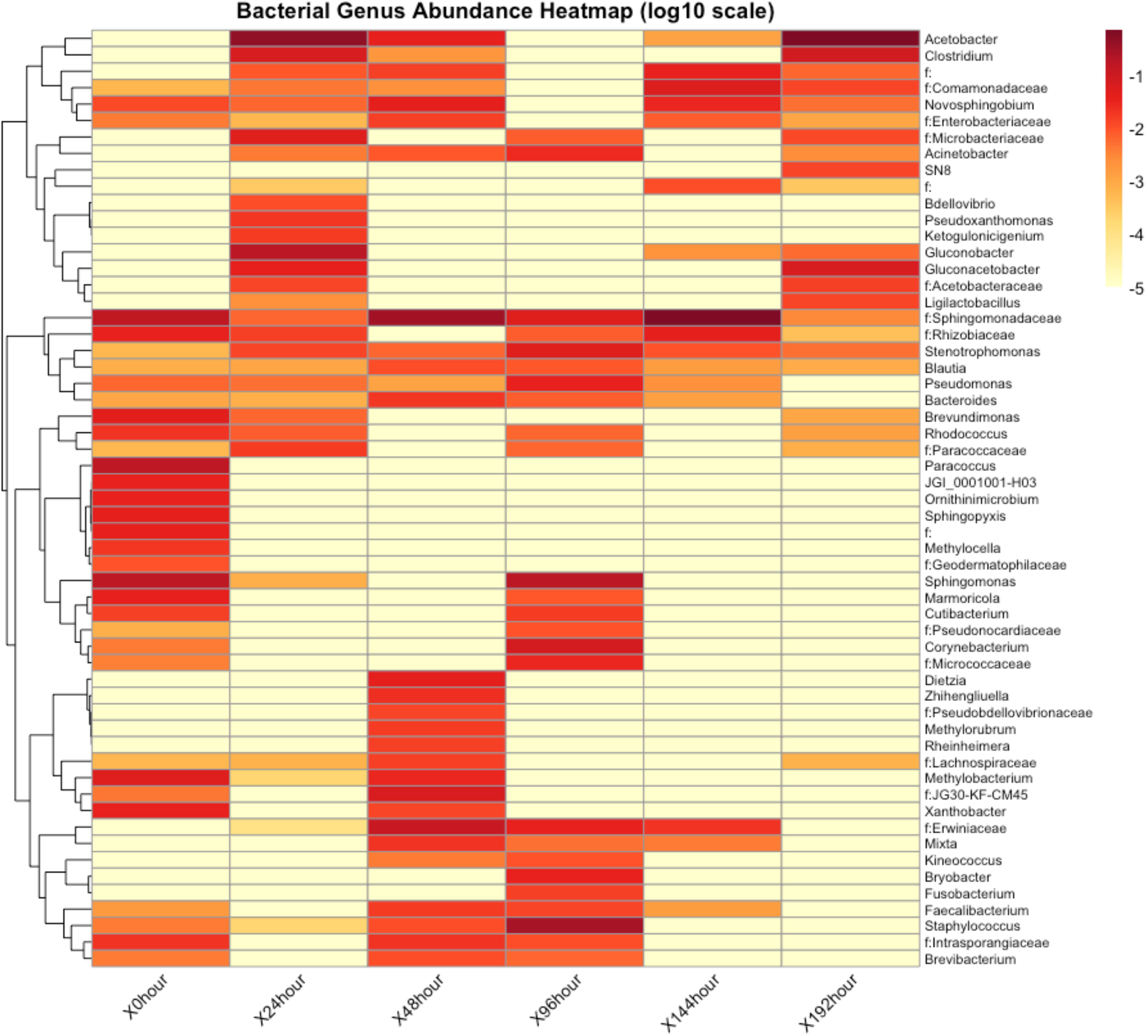
Genus-level bacterial composition during ukwa fermentation. Heatmap showing relative abundances (log10 scale) of dominant genera across time points.

### 2.6 Multivariate Analyses

Principal Component Analysis (PCA) was employed on genus-level relative abundance data to assess temporal variation in microbial community composition and its associated environmental variables. Collinearity between pH and moisture content was evaluated using Pearson correlation (r > 0.8), and highly correlated variables were excluded to minimize redundancy. Linear regression models were fitted to explore relationships among fermentation time, pH, and moisture content. Autocorrelation functions (ACF) were computed for PCA axes to evaluate temporal dependence. Redundancy Analysis (RDA) was applied to determine the influence of environmental variables and their lagged effects on microbial community structure. Ordination trajectories were used to visualize temporal shifts in microbial composition and the direction of ecological change.

## 3. RESULTS AND DISCUSSION

### 3.1 Sequencing Quality and Microbial Diversity

Microbial diversity during ukwa fermentation was characterized using high-throughput 16S rRNA sequencing across six time points (0, 24, 48, 96, 144, and 192 h). Sequencing depth ranged from approximately 37,000 to 230,000 reads per sample (median ∼75,000), with no sample dropout. Forward–reverse read concordance confirmed technical reliability (Supplementary Fig. S1A and B). Rarefaction curves plateaued at ∼5,000 reads, indicating sufficient coverage to recover core bacterial diversity (Supplementary Fig. S2; Hughes et al., 2001). ASV richness ranged from 80–95 early in fermentation but declined to ∼65–75 at later stages (Supplementary Fig. S3), reflecting the ecological narrowing commonly observed in maturing fermentations. These diversity profiles are consistent with other seed-based fermentation systems, such as Dajiang and sufu (Liang et al., 2021). Comparable diversity patterns have also been reported in African alkaline-fermented condiments such as African locust bean, further demonstrating the ecological complexity of seed-based fermentations (Parkouda et al., 2009; Obafemi et al., 2022; Adesulu-Dahunsi et al., 2022).

### 3.2 Microbial Succession, Diversity Trajectories, and Ecological Patterns

High-resolution sequencing enabled robust temporal profiling of microbial succession during ukwa fermentation. Alpha-diversity metrics showed clear restructuring of the community over time (Fig. 1). Shannon diversity exhibited temporal fluctuations, peaking at 24 h (∼4.8), declining through the mid-fermentation phase (48–96 h), and recovering to maximum values by late fermentation (192 h; ∼5.0). Simpson evenness followed a similar pattern, remaining consistently high at early (24 h) and late stages (144–192 h), indicating sustained ecological complexity despite environmental selection pressures.

Beta-diversity analysis revealed distinct temporal separation of samples, indicating dynamic community restructuring throughout fermentation (Supplementary Fig. S4). Principal component analysis explained 53% of the total variance (PC1: 29.1%, PC2: 23.9%), with samples from different fermentation stages occupying distinct regions of ordination space. The initial sample (0 h) was clearly separated from all other time points, reflecting the unique plant-associated microbiota present on unfermented seeds. Notably, early (24 h) and late (192 h) fermentation samples clustered in proximity within the upper-right quadrant, suggesting functional convergence toward aerobic, acid-producing communities at both stages. The 48 h sample occupied a distinct position in the upper-left quadrant, indicating a transient community state characterized by rapid compositional turnover. Mid-fermentation samples (96–144 h) occupied an intermediate region in ordination space, consistent with a transitional phase between early colonization and late-stage acid production. This non-linear temporal trajectory suggests that ukwa fermentation does not follow a simple unidirectional succession, but rather exhibits stage-specific community configurations driven by fluctuating environmental conditions and metabolic interactions, a pattern consistent with other spontaneous plant-based fermentations (Illeghems et al., 2013; De Vuyst & Leroy, 2020; Lavefve et al., 2021). Similar non-linear succession patterns have been reported in cocoa fermentation, where microbial community structure exhibits phase-specific clustering driven by oxygen availability, substrate depletion, and organic acid accumulation (Papalexandratou et al., 2011; De Vuyst & Leroy, 2020). The observed proximity of 24 h and 192 h samples suggests functional redundancy or convergent metabolic states at distinct fermentation stages, reflecting the complex interplay between environmental filtering and microbial niche construction (Fierer et al., 2010; Zhou & Ning, 2017).

These successional dynamics align with documented ecological patterns in cassava fermentations, where acidification and substrate depletion impose stage-specific filters on community composition (Oyewole & Odunfa, 1988; Dike et al., 2022). Similar deterministic trajectories have been reported in cocoa fermentations, where yeast, LAB, and AAB successively modify substrate chemistry and oxygen availability (Papalexandratou et al., 2011; De Vuyst & Leroy, 2020). The persistence of moderate richness at later stages suggests active niche partitioning and metabolic cross-feeding, patterns also observed in palm-sap, vegetable, and cereal-based spontaneous fermentations (Yang et al., 2020; Zhang et al., 2021; Adesulu-Dahunsi et al., 2022; Liu et al., 2025). Collectively, the distinct clustering, diversity patterns, and successional transitions indicate that ukwa fermentation follows a predictable, ecologically structured assembly process shaped by physicochemical gradients and microbial interactions.

### 3.3 Community Composition and Core Microbiome

Phylum-level profiles showed that the bacterial community was dominated by Pseudomonadota (formerly Proteobacteria) and Firmicutes (Bacillota), with a gradual shift in relative abundance toward Firmicutes as fermentation progressed. Actinomycetota declined steadily, while minor phyla remained below 10% throughout the fermentation process (Fig. 2).At the genus level (Fig. 3), the community comprised plant-associated taxa, fermentative bacteria, and acid-tolerant groups with distinct ecological functions. Genera such as Sphingomonas and Paracoccus were consistently detected and are known for their capacity to metabolize plant-derived and aromatic compounds in surface-associated environments (Asaf et al., 2020; Friedrich et al., 2008; Moungprayoon et al., 2022). Fermentative and facultatively anaerobic genera, including Clostridium and coagulase-negative Staphylococcus, were also prominent, reflecting the system’s capacity to support anaerobic metabolism and complex substrate turnover. Acidophilic taxa such as *Lactobacillus*, *Pediococcus*, *Acetobacter*, and *Gluconobacter* were likewise well represented, consistent with the acidic characteristic of mature fermentations. The core microbiome, defined as taxa present in more than 20% of samples at relative abundances above 0.01%, accounted for approximately 40% of all ASVs (Fig. 4). Members of the family Sphingomonadaceae persisted across all samples, indicating a stable ecological presence. Their persistence aligns with their documented metabolic versatility, including the ability to utilize diverse plant-derived substrates and tolerate fluctuating physicochemical conditions (Glaeser & Kämpfer, 2014; Asaf et al., 2020). Late-stage Acetobacteraceae ASVs reflected adaptation to acidic, ethanol-rich niches. The coexistence of a stable core community alongside more variable peripheral taxa is characteristic of resilient fermentation systems and provides a useful basis for future starter culture development in spontaneous fermentations (Tamang et al., 2016; Liu et al., 2023).

**Fig. 4.**
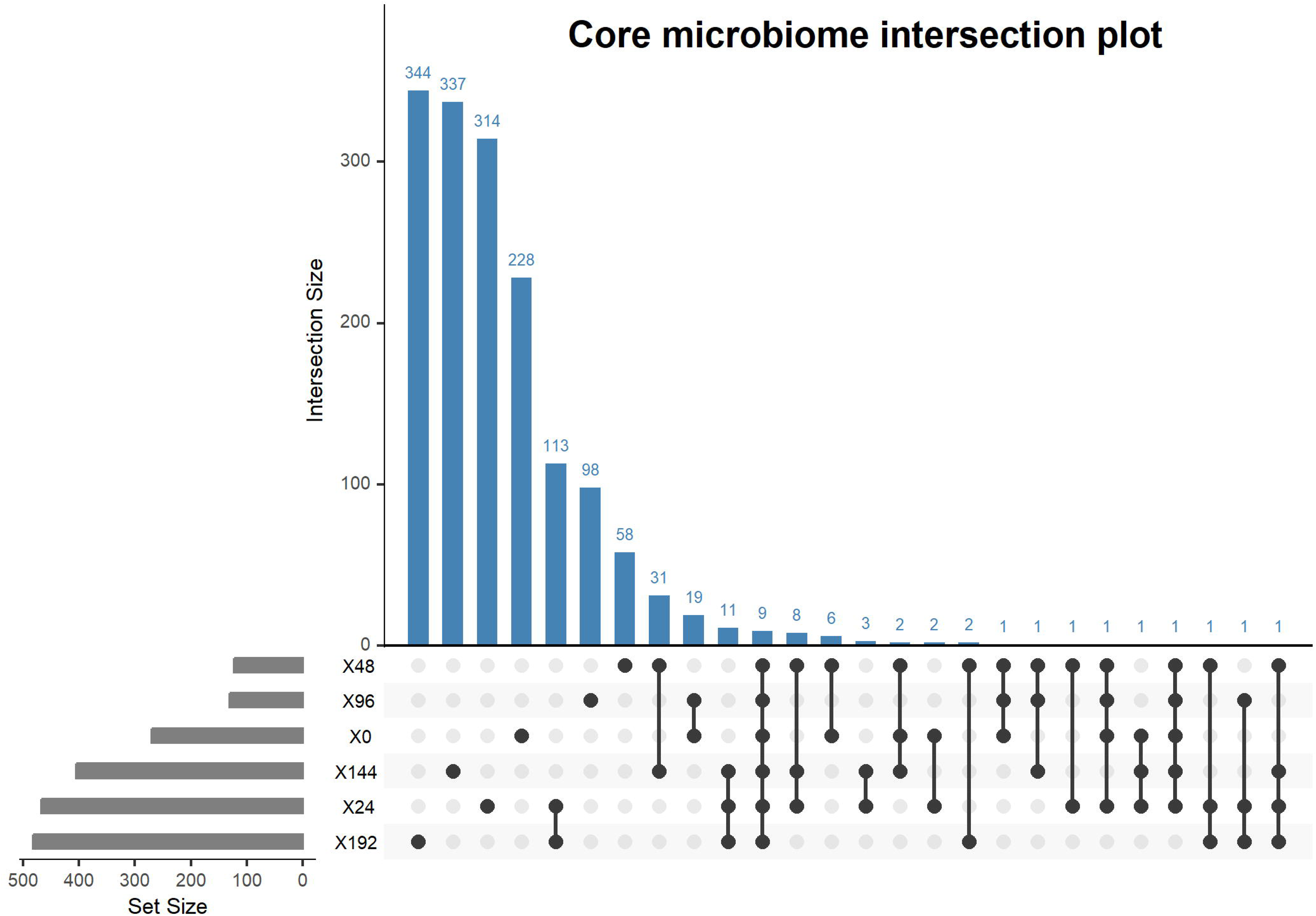
Core microbiome of ukwa fermentation. ASVs present in >20% of samples at relative abundances >0.01%.

### 3.4 Temporal Dynamics and Differential Abundance of Key Microbial Groups

Time-course patterns of the major bacterial groups revealed a clear, stage-dependent progression in community structure during ukwa fermentation (Fig. 5). In the earliest hours (0–48 h), the bacterial community showed relatively low abundances of most dominant taxa, with modest representation of Paracoccus (∼15% at 0 h), Staphylococcus (∼16% at 0 h), and low but detectable levels of Sphingomonas and Novosphingobium. These organisms are typical early colonizers of plant substrates and likely participate in the initial breakdown of surface molecules and simple carbohydrates. Members of the Erwiniaceae family (Enterobacteriaceae) showed early peak abundance at 48 h before beginning a steady decline that continued throughout fermentation. This reduction in facultative anaerobes and plant-associated taxa is likely linked to increasing acidity, reduced nutrient availability, and competitive exclusion by acid-producing bacteria (Axelsson, 2004; Tang et al., 2023). Similar declines have been reported in African cereal and root-crop fermentations as conditions become progressively more selective, often coinciding with improved microbiological safety (Adesulu-Dahunsi et al., 2022).

**Fig. 5.**
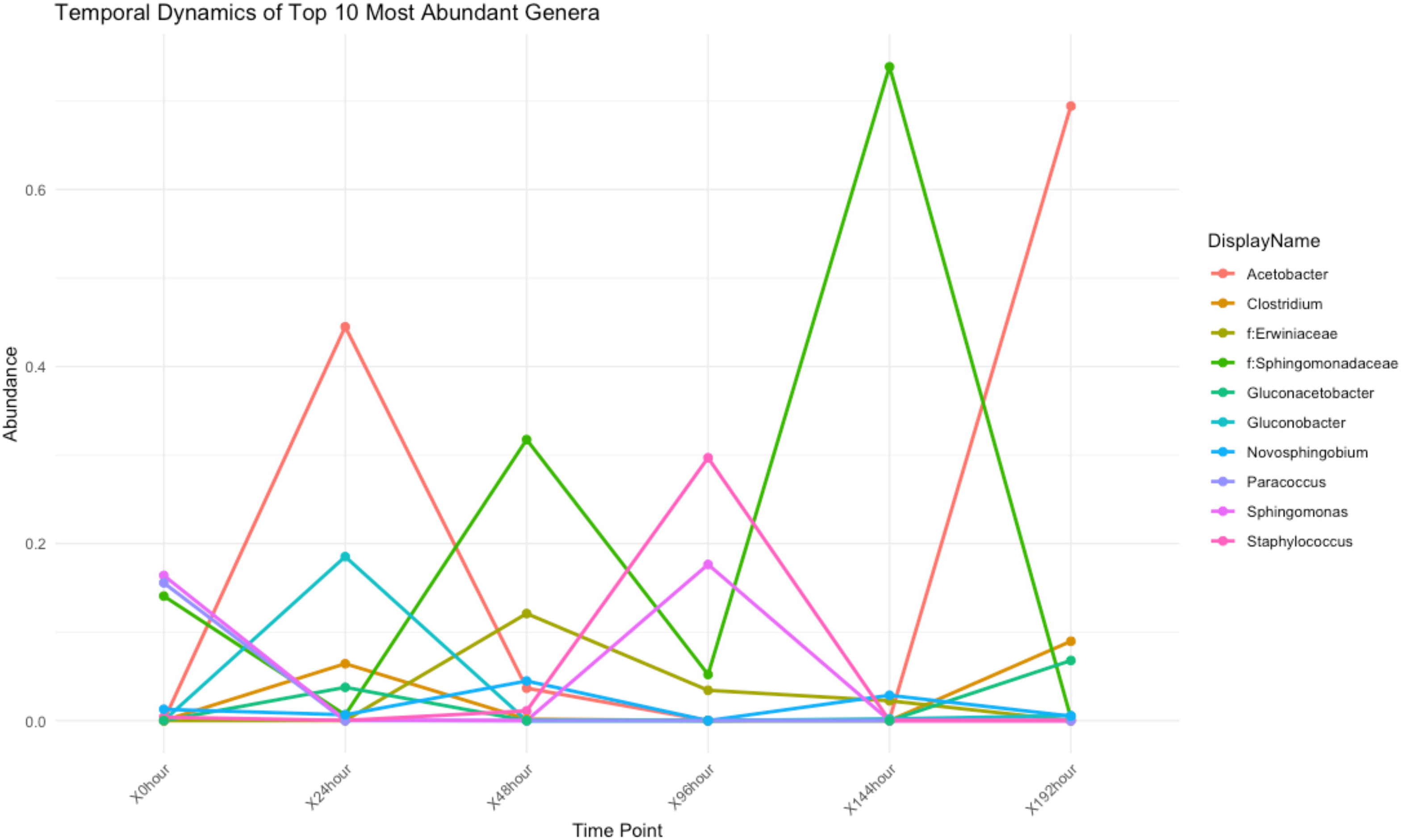
Temporal dynamics of the ten most abundant bacterial genera. Relative abundance changes across six time points (0–192 h).

As fermentation moved into the mid-phase (96–144 h), there was a dramatic shift toward specific bacterial groups. Members of the family Sphingomonadaceae increased sharply, reaching a combined peak relative abundance of approximately 80% at 144 h (Fig. 5). Concurrently, Staphylococcus showed peak abundance (∼30% at 96 h), consistent with its recognized roles in proteolysis, amino acid turnover, and flavour development in fermented foods (Heo et al., 2020). Differential abundance analysis confirmed these mid-fermentation peaks, with Staphylococcus showing maximum change at 96 h and Sphingomonadaceae at 144 h relative to the initial baseline (Fig. 8). This pattern reflects the gradual depletion of oxygen and emergence of transitional metabolic conditions. Similar patterns have been observed in cassava and other seed-based fermentations, where facultatively anaerobic taxa establish during the transition from oxygenated to oxygen-depleted conditions (Dike et al., 2022).

By the final stage of fermentation (192 h), the community shifted sharply toward acid-adapted taxa. Acetobacter became the dominant genus, reaching approximately 70% relative abundance, while Sphingomonadaceae declined dramatically (Fig. 5). Differential abundance analysis confirmed this progressive AAB enrichment, revealing a steady cumulative increase exceeding 0.6 relative to the initial baseline (Fig. 6). Other taxa including Clostridium (∼10%), members of the Erwiniaceae family, and *Gluconacetobacter* persisted at lower levels. *Gluconobacter*, *Sphingomonas*, and *Novosphingobium* remained present but at consistently low abundances throughout fermentation. This transition aligns closely with declining pH and moisture content, as well as accumulation of organic acids. The dramatic shift from Sphingomonadaceae dominance at 144 h to Acetobacter dominance at 192 h represents a functional transition from diverse aerobic metabolism to specialized acetic acid production. Comparable late-stage acetic acid bacteria dominance has been documented in cocoa, kombucha, vinegar, and several African fermented foods, where AAB prevail as the environment becomes increasingly acidic and oxidative niches re-emerge (De Vuyst & Leroy, 2020; Illeghems et al., 2013; Marsh et al., 2014; Gullo & Giudici, 2008; Tran et al., 2020; Xia et al., 2022; Díaz et al., 2019).

**Fig. 6.**
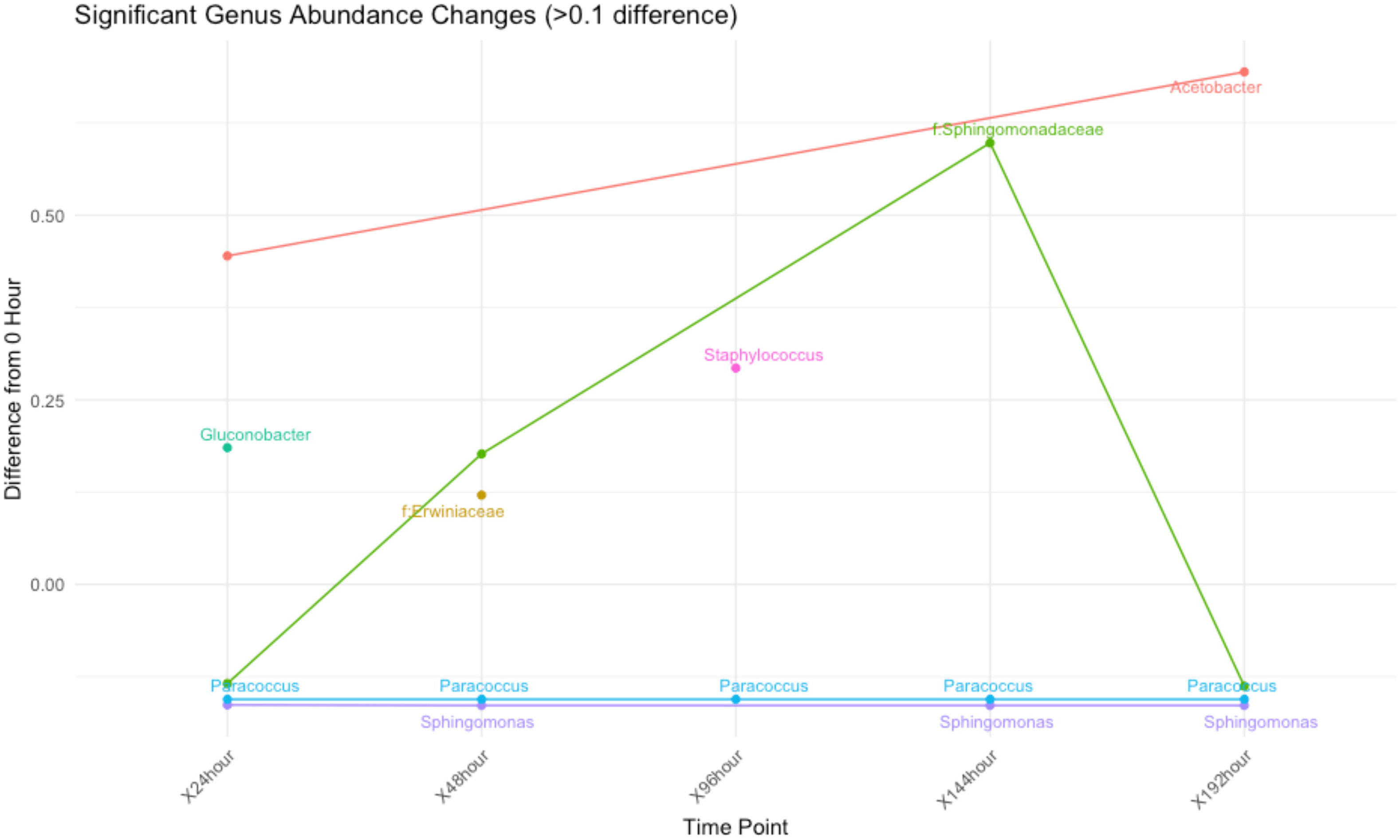
Differential abundance of bacterial genera relative to 0 h. Genera with changes >0.1 in relative abundance shown.

These temporal shifts illustrate a structured ecological sequence: early colonizers characterized by modest diversity and plant-associated taxa give way to a sharp mid-phase expansion of Sphingomonadaceae and Staphylococcus (96–144 h), which in turn is replaced by acid-tolerant acetic acid bacteria as the environment evolves toward the final stage (192 h). This stage-linked progression reflects progressive ecological filtering and is characteristic of many spontaneous fermentation systems. Comparable stage-structured restructuring has been reported in cassava fermentations (Oyewole & Odunfa, 1988) and millet-based porridges such as Hausa koko (Atter et al., 2021). Practically, this indicates that ukwa fermentation could be guided by monitoring and controlling pH and moisture trajectories. Maintaining favourable ranges may promote beneficial taxa such as Lactobacillus and acetic acid bacteria while suppressing undesirable groups. Although explicit process-control frameworks for African seed-based fermentations remain limited, similar principles underpin efforts to stabilize cereal- and root-crop fermentations. In ogi, for example, pH trajectories and titratable acidity are closely linked to LAB dominance, safety, and product quality and have been identified as key levers for process improvement and potential starter-culture standardization (Okeke et al., 2015). More broadly, reviews of African indigenous foods highlight how LAB-driven acidification and successional dynamics in cereal- and cassava-based fermentations can be exploited to enhance safety and shelf life, supporting parameter-based steering of ukwa fermentation (Adesulu-Dahunsi et al., 2022). Understanding these patterns provides a useful foundation for identifying functional indicator taxa and, ultimately, for guiding process optimization in future controlled fermentations.

### 3.5 Multivariate and Temporal Analyses

PCA (Supplementary Fig. S4) and hierarchical clustering revealed distinct temporal shifts in community structure, mirroring the time-series behavior of key microbial groups and associated environmental variables. Among these, pH and moisture (Fig. 7A) stood out as primary ecological drivers, both of which declined steadily throughout fermentation, with pH dropping at approximately 0.024 units per hour (R² = 0.97, p < 0.001) (Fig. 7B) and moisture declining proportionally (Fig. 7C). Their strong correlation (r > 0.8) reflects the kind of tightly coupled physicochemical dynamics typical of solid-state fermentation. Autocorrelation analysis of the PCA axes uncovered a pronounced temporal signature. PC1 captured stable, recurring patterns linked to dominant taxa, while PC3 reflected intermittent responses from transient groups (Supplementary Fig. S5). Correlation analysis confirmed that PC3 aligned most closely with pH gradients, while PC1 and PC5 showed weaker associations with individual physicochemical measurements (Supplementary Fig. S6). Ordination trajectory analysis (Fig. 8) revealed a non-linear path through compositional space, with samples from successive time points clearly separated along environmental gradients associated with moisture and pH decline. This directional progression reinforces the idea that these variables impose sequential constraints on microbial assembly throughout fermentation.

**Fig. 7.**
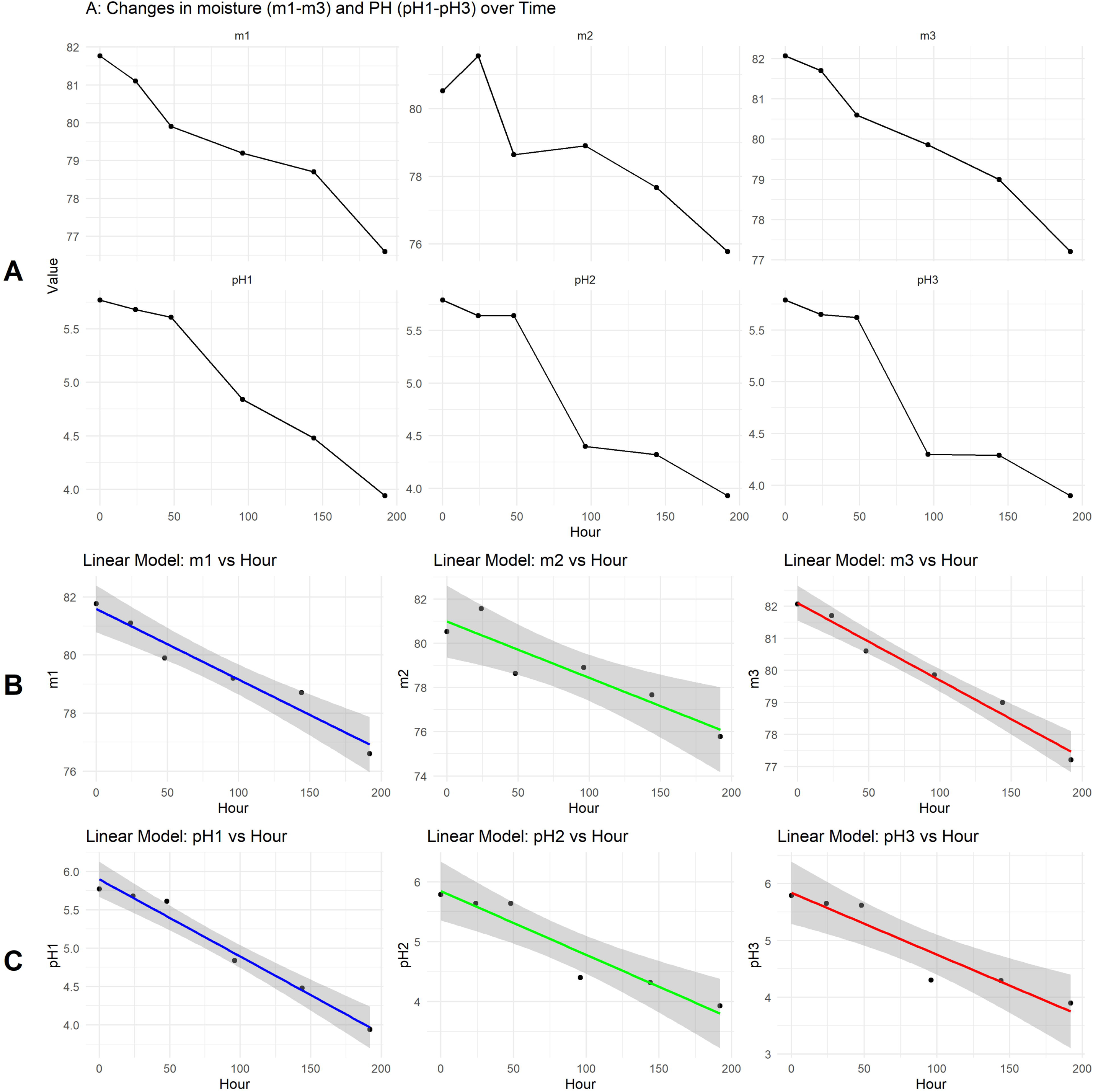
Physicochemical parameters during ukwa fermentation. (A) Temporal changes in moisture content (top: m1–m3) and pH (bottom: pH1–pH3) across three experimental replicates from 0 to 192 h. (B) Linear regression models for pH versus fermentation time in each replicate (R² > 0.97 for all replicates, p < 0.001). Average slope = 0.024 pH units per hour. (C) Linear regression models for moisture content versus fermentation time in each replicate showing consistent water loss across replicates.

RDA models further quantified these relationships (Fig. 8B). The non-lag model explained 42% of the community variation, whereas the lagged model (Fig. 8C) accounted for 79.27% of the constrained variance and exhibited stronger directional clustering (p = 0.001). This pronounced structure is consistent with an ecological memory effect, in which past environmental states influence the range of future microbial configurations.

**Fig. 8.**
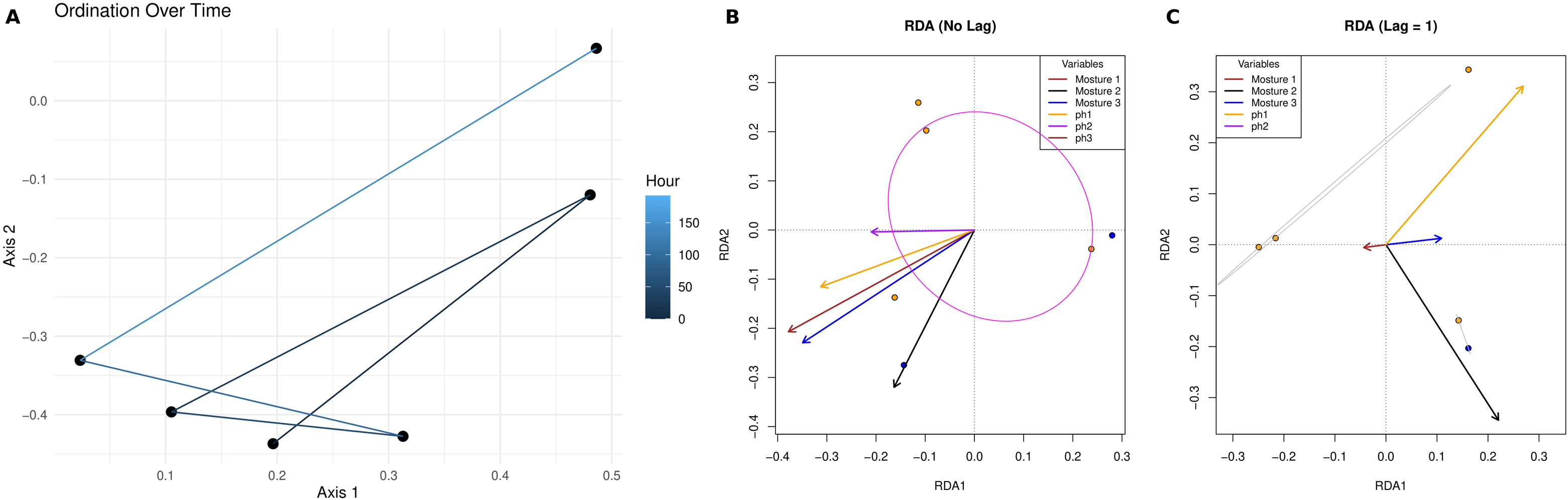
Ordination trajectory and redundancy analysis of community structure with environmental variables. (A) Temporal trajectory of bacterial community composition in ordination space (Axes 1 and 2). Points represent samples from different fermentation stages (0, 24, 48, 96, 144, and 192 h), connected by lines. Color gradient indicates fermentation time (hours). (B) Redundancy analysis (RDA) showing variation in bacterial community composition explained by concurrent pH and moisture measurements (42% of variance explained). (C) Redundancy analysis showing variation in bacterial community composition explained by one-time-step lagged pH and moisture values (79.27% of constrained variance explained, p < 0.001)

Similar forms of directional or memory-like succession have been documented in several seed-based fermented foods. In tempeh, early establishment of *Rhizopus* reshapes the matrix in ways that govern the sequence of bacterial colonization that follows (Steinkraus, 1995). Cocoa fermentations show a similarly ordered LAB–yeast–AAB progression, in which each functional group modifies the environment for the next stage of the process (De Vuyst & Leroy, 2020).

### 3.6 Co-occurrence Network Analysis and Functional Prediction

The co-occurrence network resolved six distinct modules (M1–M6) characterized by strong positive correlations (r = 0.98–1.00), indicating that the bacterial community is organized around interacting microbial guilds rather than isolated taxa (Fig. 9A). Module M1, enriched with *Weissella*, *Bdellovibrio*, and *Corynebacterium*, was associated with early carbohydrate fermentation and rapid substrate turnover. Comparable early-stage communities, dominated by heterofermentative lactic acid bacteria, have been described in vegetable and cereal fermentations, such as kimchi and sourdough, where these organisms help initiate fermentation and shape early community development (Kim et al., 2016; Fu et al., 2022).

**Fig. 9.**
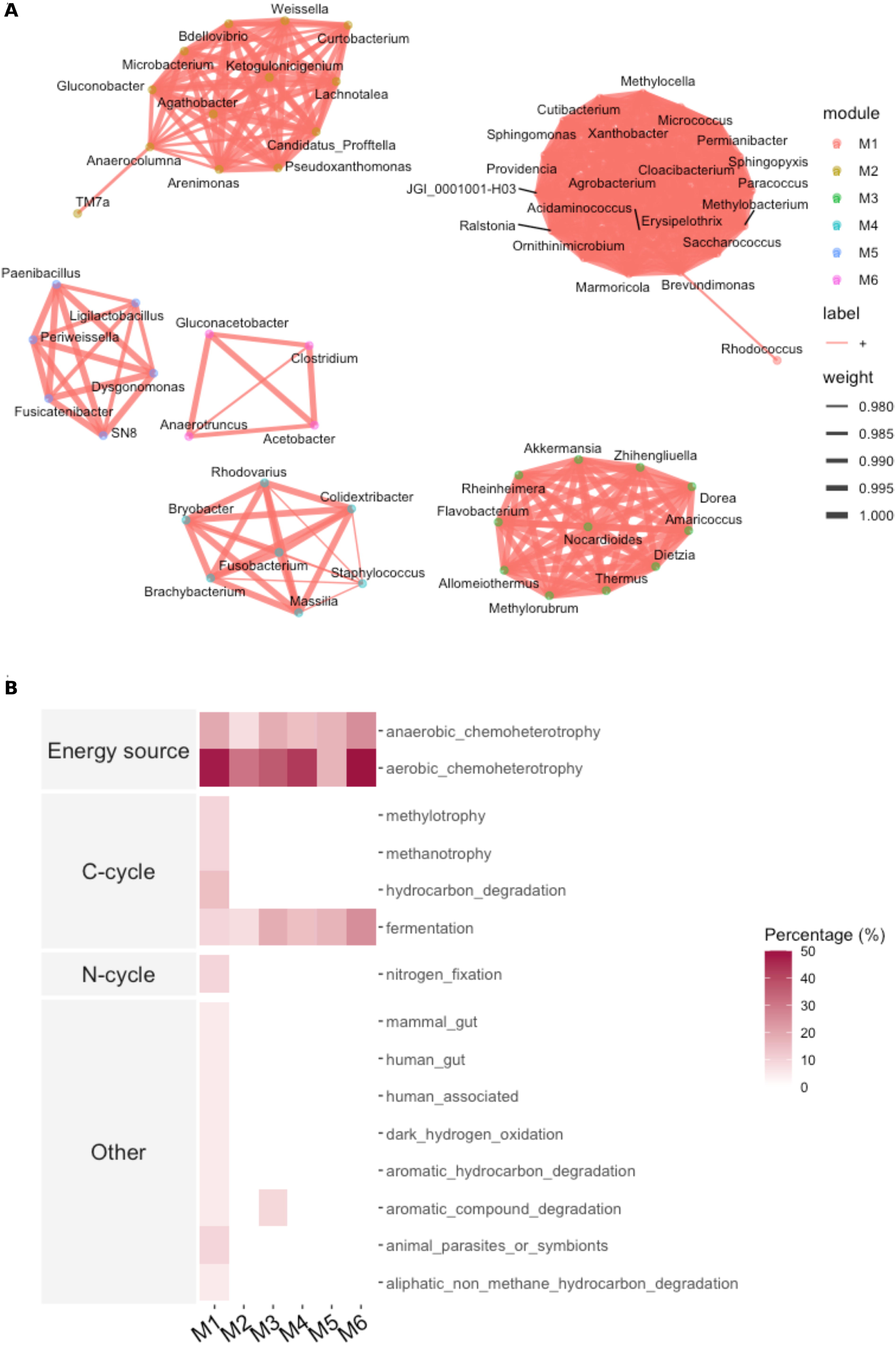
Co-occurrence network and functional predictions. (A) Network showing modules M1–M6 with significant correlations (r = 0.98–1.00). (B) FAPROTAX-predicted functional profiles by module.

Module M3, containing *Providencia* and *Erysipelothrix*, appeared linked to amino acid and peptide turnover. In contrast, Module M4, comprising *Akkermansia*, Rheinheimera, and Nocardioides, represented late-phase taxa adapted to lower pH and reduced moisture, supporting polysaccharide degradation, stress tolerance, and potential cross-feeding interactions. Modules M5 and M6 were smaller but strongly interconnected, suggesting narrow yet coordinated metabolic roles. Comparable modular structures have been reported in fermented food ecosystems and are considered hallmarks of community resilience and functional redundancy (Huang et al., 2024; Liu et al., 2023).

Functional annotation using FAPROTAX revealed the predicted metabolic potentials associated with each module (Fig. 9B). As FAPROTAX assignments are based on trait information from characterized taxa, these results reflect predicted metabolic capabilities rather than directly measured functions. Under this framework, Modules M1, M2, and M4 were enriched in chemoheterotrophic and carbohydrate-utilization pathways, consistent with their dominant genera and prevalence during active fermentation stages. In contrast, Modules M3 and M6 were associated with functions related to aromatic compound degradation and broader carbon-cycle processes, while Module M5 included nitrogen-cycle traits such as nitrate reduction.

These patterns indicate that the bacterial community is structured into complementary functional guilds operating across successive phases of fermentation. Environmental gradients described in Section 3.4 further shaped module distribution, with early modules favoured under higher moisture and moderate acidity, and late modules enriched in acid-tolerant taxa. This environmentally driven structuring aligns with deterministic succession commonly observed in spontaneous fermentations (Gautam et al., 2025). Overall, the network topology, temporal dynamics, and predicted functional profiles support interpreting ukwa fermentation as a self-organizing microbial consortium in which interacting modules sustain metabolic continuity, stability, and ecological coherence throughout the fermentation process (Liu et al., 2023).

### 3.7 Insights and Implications for ukwa Fermentation and African Microbial Ecology

The results of this study deepen current understanding of ukwa fermentation and contribute to the broader knowledge of African fermented food systems. By resolving successional trajectories, distinguishing core and transient taxa, and linking these patterns to environmental drivers, this work provides a foundation for improving process consistency and product quality. From an applied perspective, a phased starter culture strategy emerges as a promising avenue. Such an approach could involve early inoculation with epiphytic substrate degraders, mid-phase reinforcement with key Firmicutes, and late-phase dominance by acid- and oxidation-tolerant taxa such as Lactobacillus and Acetobacter, reflecting successional dynamics observed in toddy and other spontaneous fermentations (Das & Tamang, 2021). While African fermentations are often described as predominantly LAB-driven and relatively stable (Adesulu-Dahunsi et al., 2022), the present findings reveal a more dynamic, multi-phase succession in seed-based systems such as ukwa. More broadly, these results underscore the importance of region-specific microbial ecology studies and illustrate how traditional fermentations can provide insight into microbial function, interaction, and community assembly.

### 3.8 Limitations and Future Research Directions

This work provides the first next-generation sequencing profile of ukwa fermentation and establishes a basis for future investigation. While pH and moisture were monitored as key environmental parameters, extending physicochemical characterization to include redox potential, oxygen dynamics, and temperature gradients may reveal additional ecological transitions. Increased sampling frequency could also capture transient metabolic states occurring between the current time points.

Although this study was based on a single traditionally fermented batch, such a design is common in first-generation microbiome studies of under-characterized food systems, where establishing temporal baselines is a necessary precursor to replicated, hypothesis-driven work. The strong successional structure, high sequencing depth, and clear physicochemical gradients observed here provide substantial internal robustness for interpreting ecological patterns. Future studies incorporating replicated fermentations across processors, seasons, and regions will be essential for evaluating process variability and refining ecological models of ukwa fermentation. Integrating metabolomics, metatranscriptomics, volatilomics, or shotgun metagenomics will further clarify the functional consequences of taxonomic shifts and their implications for flavour development, safety, and nutritional quality (Tamang et al., 2020).

## 4. CONCLUSION

This study presents the first high-throughput sequencing analysis of microbial succession during spontaneous ukwa (Treculia africana) fermentation and establishes a foundational ecological framework for this under-characterized African food system. Ukwa fermentation follows a structured, stage-wise trajectory, beginning with epiphytic, oxygen-tolerant taxa, transitioning through a mid-phase dominated by fermentative Firmicutes, and culminating in late-stage acid-and oxidation-tolerant specialists, including lactic and acetic acid bacteria. These transitions closely tracked declines in pH and moisture and were accompanied by distinct co-occurrence modules and a persistent core microbiome.

The ecological patterns identified here extend broader understanding of microbial assembly in solid-state fermentations and highlight the role of environmental filtering in shaping community transitions. By defining key taxa, functional guilds, and physicochemical drivers, this study provides a mechanistic basis for improving process control, guiding starter culture development, and enhancing the reproducibility, safety, and sensory quality of this culturally significant traditional food.

## 5. PRACTICAL APPLICATION

Elucidating the microbial succession and functional organization underlying ukwa fermentation provides a practical foundation for strengthening traditional processing methods. Identifying stage-specific taxa and their environmental drivers enables the rational design of starter cultures that replicate desirable natural fermentation trajectories. The predictable relationship between pH decline and microbial succession (R² = 0.97) offers a straightforward approach for monitoring fermentation progress in small-scale production settings, where pH measurement is simple and affordable. The core taxa identified here—particularly members of Sphingomonadaceae and Acetobacteraceae that persisted across fermentation stages, along with Lactobacillaceae that drive mid-to-late-stage acidification, represent promising candidates for developing functional starter cultures tailored to ukwa fermentation. Integrating these insights into artisanal or semi-controlled production systems can enhance product consistency, safety, and flavour quality while preserving the cultural authenticity, nutritional value, and local relevance of this indigenous African food. Controlled fermentation trials evaluating these core organisms as starter culture components will be needed to translate these ecological findings into practical processing improvements.

## 6. ACKNOWLEDGMENTS

This research received no specific grant from any funding agency. The authors acknowledge Dr. Muiz O. Akinyemi (Leeds Institute of Health Sciences, University of Leeds, UK) for assistance with initial bioinformatics analysis and Laragen Inc. (Culver City, CA, USA) for sequencing and bioinformatics services.

## 8. DATA AVAILABILITY

The raw 16S rRNA gene sequencing reads generated in this study have been deposited in the NCBI Sequence Read Archive under BioProject accession number PRJNA681567. The ukwa fermentation samples analyzed in this manuscript correspond to samples within this BioProject. Cassava fermentation data from the same sequencing project were reported separately (Dike et al., 2022).

## 9. CRediT AUTHORSHIP CONTRIBUTION STATEMENT

**Kelechi Stanley Dike**: Conceptualization, Methodology, Investigation, Validation, Formal analysis, Writing – original draft, Writing – review & editing. **Chidinma Francisca Ezeh**: Methodology, Investigation, Formal analysis. **Monicah Maseke**: Data curation, Software, Formal analysis, Writing – review & editing. **Chinwe Happiness Justice-Alucho:** Methodology, Writing – review & editing. **Chibundu N. Ezekiel**: Resources, Writing – review & editing.

## REFERENCES

Achi, O.K., 1992. Microorganisms associated with natural fermentation of Prosopis africana seeds for the production of okpiye. Plant Foods Hum. Nutr. 42, 297–304. 10.1007/BF02194090

Adesulu-Dahunsi, A.T., Jeyaram, K., Sanni, A.I., 2022. Co-occurrence of Lactobacillus species during fermentation of African indigenous foods: impact on food safety and shelf-life extension. Front. Microbiol. 13, 684730. 10.3389/fmicb.2022.684730

Andrews, S., 2010. FastQC: A Quality Control Tool for High Throughput Sequence Data. Babraham Bioinformatics, Babraham Institute, Cambridge, United Kingdom. https://www.bioinformatics.babraham.ac.uk/projects/fastqc

Asaf, S., Numan, M., Khan, A.L., Al-Harrasi, A., 2020. Sphingomonas: from diversity and genomics to functional role in environmental remediation and plant growth. Crit. Rev. Biotechnol. 40, 138–152. 10.1080/07388551.2019.1709793

Atter, A., Diaz, M., Kunadu, A.P., Mayer, M.J., Colquhoun, I.J., Nielsen, D.S., Baker, D., Narbad, A., 2021. Microbial diversity and metabolite profile of fermenting millet in the production of Hausa koko, a Ghanaian fermented cereal porridge. Front. Microbiol. 12, 681983. 10.3389/fmicb.2021.681983

Auchtung, T.A., Hallen-Adams, H.E., Hutkins, R.W., 2025. Microbial interactions and ecology in fermented food ecosystems. Nat. Rev. Microbiol. 23, 622–634. 10.1038/s41579-025-01191-w

Axelsson, L., 2004. Lactic acid bacteria: classification and physiology. In: Salminen, S., von Wright, A., Ouwehand, A. (Eds.), Lactic Acid Bacteria: Microbiological and Functional Aspects, third ed. Marcel Dekker, New York, pp. 1–66.

Azokpota, P., Hounhouigan, D.J., Nago, M.C., 2006. Microbiological and chemical changes during the fermentation of African locust bean (Parkia biglobosa) to produce afitin, iru, and sonru. Int. J. Food Microbiol. 107, 304–309. 10.1016/j.ijfoodmicro.2005.10.026

Bastian, M., Heymann, S., Jacomy, M., 2009. Gephi: an open source software for exploring and manipulating networks. Proc. Int. AAAI Conf. Weblogs Soc. Media 3, 361–362.

Bolyen, E., Rideout, J.R., Dillon, M.R., et al., 2019. Reproducible, interactive, scalable and extensible microbiome data science using QIIME 2. Nat. Biotechnol. 37, 852–857. 10.1038/s41587-019-0209-9

Callahan, B.J., McMurdie, P.J., Rosen, M.J., et al., 2016. DADA2: high-resolution sample inference from Illumina amplicon data. Nat. Methods 13, 581–583. 10.1038/nmeth.3869

Csárdi, G., Nepusz, T., 2006. The igraph software package for complex network research. InterJournal Complex Syst. 1695, 1–9.

Das, S., Tamang, J.P., 2021. Changes in microbial communities and their predictive functionalities during fermentation of toddy, an alcoholic beverage of India. Microbiol. Res. 248, 126769. 10.1016/j.micres.2021.126769

De Filippis, F., Parente, E., Ercolini, D., 2017. Metagenomics insights into food fermentations. Microb. Biotechnol. 10, 91–102. 10.1111/1751-7915.12421

De Filippis, F., Troise, A.D., Vitaglione, P., Ercolini, D., 2018a. Different temperatures select distinctive acetic acid bacteria species and promote organic acids production during Kombucha tea fermentation. Food Microbiol. 73, 11–16. 10.1016/j.fm.2018.01.008

De Filippis, F., Parente, E., Ercolini, D., 2018b. Recent past, present, and future of the food microbiome. Annu. Rev. Food Sci. Technol. 9, 589–608. 10.1146/annurev-food-030117-012312

De Vuyst, L., Leroy, F., 2020. Functional role of yeasts, lactic acid bacteria and acetic acid bacteria in cocoa fermentation processes. FEMS Microbiol. Rev. 44, 432–453. 10.1093/femsre/fuaa014

Díaz, M., Ladero, V., del Rio, B., Redruello, B., Fernández, M., Alvarez, M.A., 2019. Comparison of the microbial composition of African fermented foods using amplicon sequencing. Sci. Rep. 9, 13863. 10.1038/s41598-019-50190-4

Dike, K.S., 2024. Analysis of the bacterial diversity during the fermentation of ukwa (Treculia africana Decne). World News Nat. Sci. 57, 59–67.

Dike, K.S., Okafor, C.P., Ohabughiro, B.N., Maduwuba, M.C., Ezeokoli, O.T., Ayeni, K.I., Okafor, C.M., Ezekiel, C.N., 2022. Analysis of bacterial communities of three cassava-based traditionally fermented Nigerian foods. Lett. Appl. Microbiol. 74, 452–460. 10.1111/lam.13621

Ercolini, D., 2017. High-throughput sequencing and microbial food ecology. Curr. Opin. Food Sci. 14, 76–82. 10.1016/j.cofs.2017.07.001

Fierer, N., Nemergut, D., Knight, R., Craine, J.M., 2010. Changes through time: integrating microorganisms into the study of succession. Res. Microbiol. 161, 635–642. 10.1016/j.resmic.2010.06.002

Friedrich, C.G., Quentmeier, A., Bardischewsky, F., Rother, D., Orawski, G., Hellwig, P., Fischer, J., 2008. Redox control of chemotrophic sulfur oxidation of Paracoccus pantotrophus. In: Dahl, C., Friedrich, C.G. (Eds.), Microbial Sulfur Metabolism. Springer, Berlin, Heidelberg, pp. 139–150. 10.1007/978-3-540-72682-1_12

Fu, L., Zhao, C., Chen, H., Lu, W., Lv, M., 2022. Relationship between microbial composition of sourdough and its physicochemical properties. Foods 11, 1924. 10.3390/foods11131924

Gautam, A., Poopalarajah, R., Ahmad, A.R., et al., 2025. Ecological factors that drive microbial communities in culturally diverse fermented foods. BMC Microbiol. 25, 655. 10.1186/s12866-025-04413-6

Glaeser, S.P., Kämpfer, P., 2014. The family Sphingomonadaceae. In: Rosenberg, E., DeLong, E.F., Lory, S., Stackebrandt, E., Thompson, F. (Eds.), The Prokaryotes: Alphaproteobacteria and Betaproteobacteria. Springer, Berlin, Heidelberg, pp. 641–707. 10.1007/978-3-642-30197-1_302

Gloor, G.B., Macklaim, J.M., Pawlowsky-Glahn, V., Egozcue, J.J., 2017. Microbiome datasets are compositional: and this is not optional. Front. Microbiol. 8, 2224. 10.3389/fmicb.2017.02224

Gullo, M., Giudici, P., 2008. Acetic acid bacteria in traditional balsamic vinegar: phenotypic traits relevant for starter culture selection. Int. J. Food Microbiol. 125, 46–53. 10.1016/j.ijfoodmicro.2007.11.076

Hao, F., Tan, Y., Lv, X., Liu, Y., Li, Y., 2021. Microbial community succession and environmental driving factors during initial fermentation of Maotai-flavor Baijiu. Front. Microbiol. 12, 669201. 10.3389/fmicb.2021.669201

Heo, S., Lee, J.H., Jeong, D.W., 2020. Food-derived coagulase-negative Staphylococcus as starter cultures for fermented foods. Food Sci. Biotechnol. 29, 1023–1035. 10.1007/s10068-020-00789-5

Hill, L.K., Gupta, A.K., Sharma, M., Jha, A.K., 2025. A comparative review of breadfruit seeds (Treculia africana, Artocarpus nobilis, and Artocarpus heterophyllus): nutritional composition, bioactive compounds, and processing techniques. S. Afr. J. Bot. 164, 12–25. 10.1016/j.sajb.2025.03.062

Huang, Z., Zeng, B., Deng, J., et al., 2024. Succession of microbial community structure in fermented grains during strong-flavor Baijiu fermentation and its impact on the metabolism of acids, alcohols, and esters. Food Sci. Biotechnol. 10.1007/s10068-024-01591-3

Hughes, J.B., Hellmann, J.J., Ricketts, T.H., Bohannan, B.J.M., 2001. Counting the uncountable: statistical approaches to estimating microbial diversity. Appl. Environ. Microbiol. 67, 4399–4406. 10.1128/AEM.67.10.4399-4406.2001

Illeghems, K., De Vuyst, L., Weckx, S., 2013. Complete genome sequence and comparative analysis of Acetobacter pasteurianus 386B, a strain well-adapted to the cocoa bean fermentation ecosystem. BMC Genomics 14, 526. 10.1186/1471-2164-14-526

Kim, H.Y., Kim, J., Han, S.K., 2016. Heterofermentative lactic acid bacteria dominate in Korean commercial kimchi. J. Microbiol. Biotechnol. 26, 1493–1500. 10.4014/jmb.1603.03012

Klindworth, A., Pruesse, E., Schweer, T., Peplies, J., Quast, C., Horn, M., Glöckner, F.O., 2013. Evaluation of general 16S ribosomal RNA gene PCR primers for classical and next-generation sequencing-based diversity studies. Nucleic Acids Res. 41, e1. 10.1093/nar/gks808

Lavefve, L., Cureau, N., Rodhouse, L., Marasini, D., Walker, L.M., Ashley, D., Lee, S.O., Gadonna-Widehem, P., Anton, P.M., Carbonero, F., 2021. Microbiota profiles and dynamics in fermented plant-based products and preliminary assessment of their in vitro gut microbiota modulation. Food Front. 2, 268–281. 10.1002/fft2.75

Leroy, F., De Vuyst, L., 2004. Lactic acid bacteria as potential functional starter cultures for the food fermentation industry. Trends Food Sci. Technol. 15, 67–78. 10.1016/j.tifs.2003.09.004

Liang, T., Xie, X., Ma, J., Wu, L., Xi, Y., Zhao, H., Li, L., Li, H., Feng, Y., Xue, L., Chen, M., Chen, X., Zhang, J., Ding, Y., Wu, Q., 2021. Microbial communities and physicochemical characteristics of traditional Dajiang and sufu in North China revealed by high-throughput sequencing of 16S rRNA. Front. Microbiol. 12, 665243. 10.3389/fmicb.2021.665243

Liu, W., Tang, Y., Zhang, J., Bai, J., Zhu, Y., Zhu, L., Zhao, Y., Daglia, M., Xiao, X., He, Y., 2025. Microbial interactions in food fermentation: interactions, analysis strategies, and quality enhancement. Foods 14, 2515. 10.3390/foods14142515

Liu, Y., Pan, Y., Zhang, Y., Wang, X., 2023. Integrated meta-omics approaches reveal Saccharopolyspora as the core functional genus in Chinese Daqu fermentation. NPJ Biofilms Microbiomes 9, 32. 10.1038/s41522-023-00432-1

Louca, S., Parfrey, L.W., Doebeli, M., 2016. Decoupling function and taxonomy in the global ocean microbiome. Science 353, 1272–1277. 10.1126/science.aaf4507

Marsh, A.J., O’Sullivan, O., Hill, C., Ross, R.P., Cotter, P.D., 2014. Sequence-based analysis of the bacterial and fungal compositions of multiple kombucha (tea fungus) samples. Food Microbiol. 38, 171–178. 10.1016/j.fm.2013.09.003

McMurdie, P.J., Holmes, S., 2013. phyloseq: an R package for reproducible interactive analysis and graphics of microbiome census data. PLoS One 8, e61217. 10.1371/journal.pone.0061217

McMurdie, P.J., Holmes, S., 2014. Waste not, want not: why rarefying microbiome data is inadmissible. PLoS Comput. Biol. 10, e1003531. 10.1371/journal.pcbi.1003531

Moungprayoon, A., Lunprom, S., Reungsang, A., Salakkam, A., 2022. High cell-density cultivation of Paracoccus sp. on sugarcane juice for poly(3-hydroxybutyrate) production. Front. Bioeng. Biotechnol. 10, 878688. 10.3389/fbioe.2022.878688

Obafemi, Y.D., Oranusi, S., Omemu, A.M., et al., 2022. African fermented foods: overview, emerging benefits, and novel approaches to microbiome profiling. NPJ Sci. Food 6, 20. 10.1038/s41538-022-00130-w

Ojimelukwe, P.C., Ugwuona, F.U., 2021. The traditional and medicinal use of African breadfruit (Treculia africana Decne): an underutilized ethnic food of the Ibo tribe of South East, Nigeria. J. Ethn. Foods 8, 35. 10.1186/s42779-021-00097-1

Okeke, C.A., Ezekiel, C.N., Nwangburuka, C.C., Sulyok, M., Ezeamagu, C.O., Adeleke, R.A., Dike, S.K., Krska, R., 2015. Bacterial diversity and mycotoxin reduction during maize fermentation (steeping) for ogi production. Front. Microbiol. 6, 1402. 10.3389/fmicb.2015.01402

Ouoba, L.I., Diawara, B., Annan, N.T., Poll, L., Jakobsen, M., 2005. Volatile compounds of Soumbala, a fermented African locust bean (Parkia biglobosa) food condiment. J. Appl. Microbiol. 99, 1413–1421. 10.1111/j.1365-2672.2005.02722.x

Oyewole, O.B., Odunfa, S.A., 1988. Microbiological studies on cassava fermentation for lafun production. Food Microbiol. 5, 125–133. 10.1016/0740-0020(88)90012-9

Oyetayo, F.L., Oyetayo, V.O., 2020. The African breadfruit (Treculia africana Decne) plant seed: a potential source of essential food and medicinal phytoconstituents. In: Preedy, V.R., Watson, R.R. (Eds.), Nuts and Seeds in Health and Disease Prevention, second ed. Academic Press, Cambridge, MA, pp. 45–50. 10.1016/B978-0-12-818553-7.00004-8

Pandey, A., Soccol, C.R., Mitchell, D., 2000. New developments in solid-state fermentation: bioprocesses and products. Process Biochem. 35, 1153–1169. 10.1016/S0032-9592(00)00152-7

Papalexandratou, Z., Camu, N., Falony, G., De Vuyst, L., 2011. Comparison of the bacterial species diversity of spontaneous cocoa bean fermentations carried out at selected farms in Ivory Coast and Brazil. Food Microbiol. 28, 964–973. 10.1016/j.fm.2011.01.010

Parkouda, C., Nielsen, D.S., Azokpota, P., Annan, N.T., Møller, P.L., Amoa-Awua, W.K., Hounhouigan, D.J., Jakobsen, M., 2009. The microbiology of alkaline-fermentation of indigenous seeds used as food condiments in Africa and Asia. Crit. Rev. Microbiol. 35, 139–156. 10.1080/10408410902793056

Peschel, S., Müller, C.L., von Mutius, E., Boulesteix, A.L., Depner, M., 2021. NetCoMi: network construction and comparison for microbiome data in R. Brief. Bioinform. 22, bbaa290. 10.1093/bib/bbaa290

Quast, C., Pruesse, E., Yilmaz, P., Gerken, J., Schweer, T., Yarza, P., Peplies, J., Glöckner, F.O., 2013. The SILVA ribosomal RNA gene database project: improved data processing and web-based tools. Nucleic Acids Res. 41, D590–D596. 10.1093/nar/gks1219

R Core Team, 2024. R: A Language and Environment for Statistical Computing. R Foundation for Statistical Computing, Vienna, Austria. https://www.R-project.org/

Schloss, P.D., 2024. Rarefaction is currently the best approach to control for uneven sequencing effort in amplicon sequence analyses. mSphere 9, e00354–23. 10.1128/msphere.00354-23

Segata, N., Izard, J., Waldron, L., Gevers, D., Miropolsky, L., Garrett, W.S., Huttenhower, C., 2011. Metagenomic biomarker discovery and explanation. Genome Biol. 12, R60. 10.1186/gb-2011-12-6-r60

Steinkraus, K.H., 1995. Handbook of Indigenous Fermented Foods, second ed. CRC Press, Boca Raton, FL.

Tamang, J.P., Shin, D.H., Jung, S.J., Chae, S.W., 2016. Functional properties of microorganisms in fermented foods. Front. Microbiol. 7, 578. 10.3389/fmicb.2016.00578

Tamang, J.P., Cotter, P.D., Endo, A., et al., 2020. Fermented foods in global health: microbiome interactions and nutritional impact. Compr. Rev. Food Sci. Food Saf. 19, 184–217. 10.1111/1541-4337.12520

Tang, H., Huang, W., Yao, Y.F., 2023. The metabolites of lactic acid bacteria: classification, biosynthesis and modulation of gut microbiota. Microb. Cell 10, 49–62. 10.15698/mic2023.03.792

Tran, T., Grandvalet, C., Verdier, F., Martin, A., Alexandre, H., Tourdot-Maréchal, R., 2020. Microbial dynamics between yeasts and acetic acid bacteria in Kombucha: impacts on the chemical composition of the beverage. Foods 9, 963. 10.3390/foods9070963

Xia, M., Zhang, X., Xiao, Y., Sheng, Q., Tu, L., Chen, F., Yan, Y., Zheng, Y., Wang, M., 2022. Interaction of acetic acid bacteria and lactic acid bacteria in multispecies solid-state fermentation of traditional Chinese cereal vinegar. Front. Microbiol. 13, 964855. 10.3389/fmicb.2022.964855

Yan, Y., Sun, Y., Cui, J., Zhang, Y., Lv, Z., 2025. Environmental factors and microbial interactions drive microbial community succession during solid-state fermentation of corn husk for microbial biomass protein production. Front. Microbiol. 16, 1646555. 10.3389/fmicb.2025.1646555

Yang, X., Hu, W., Xiu, Z., Jiang, A., Yang, X., Saren, G., Ji, Y., Guan, Y., Feng, K., 2020. Microbial community dynamics and metabolome changes during spontaneous fermentation of Northeast sauerkraut from different households. Front. Microbiol. 11, 1878. 10.3389/fmicb.2020.01878

Zhao, Y., Feng, X., Zhang, X., Li, H., Wang, Y., Wang, X., 2023. Microbial community succession and its correlation with quality characteristics during gray sufu fermentation. Foods 12, 2767. 10.3390/foods12142767

Zhou, J., Ning, D., 2017. Stochastic community assembly: does it matter in microbial ecology? Microbiol. Mol. Biol. Rev. 81, e00002–17. 10.1128/MMBR.00002-17

