## Supplementary figures and images for "Bacterial succession, functional organization, and ecological drivers during spontaneous ukwa (*Treculia africana*) fermentation"

### Supplementary Fig 1

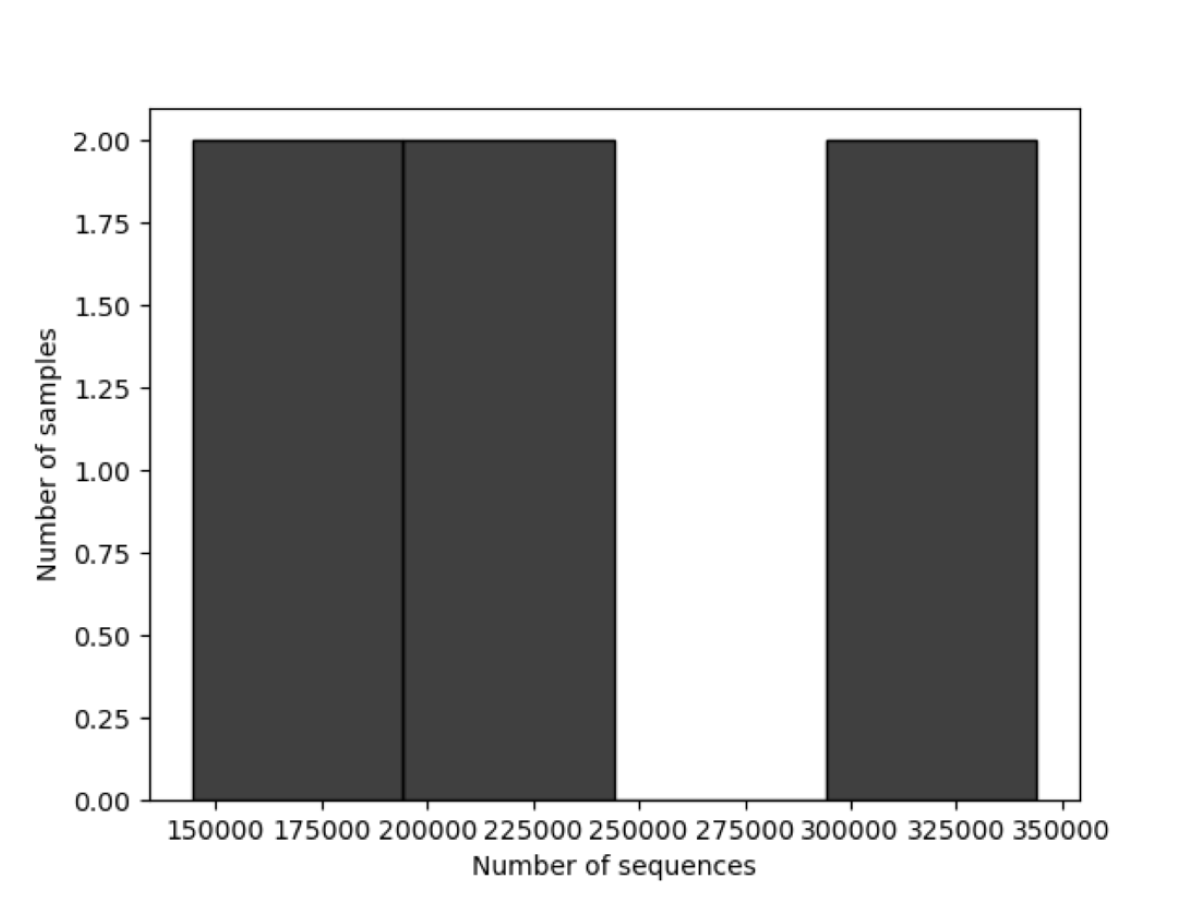

### Supplementary Fig 2

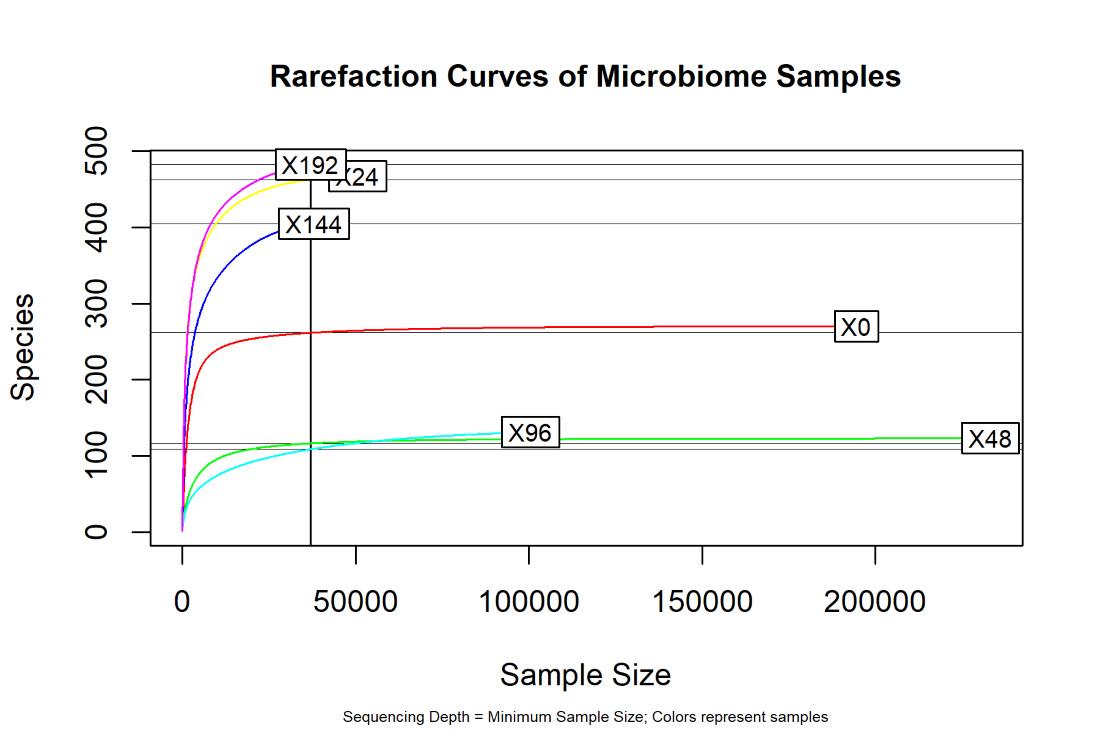

### Supplementary Fig 3

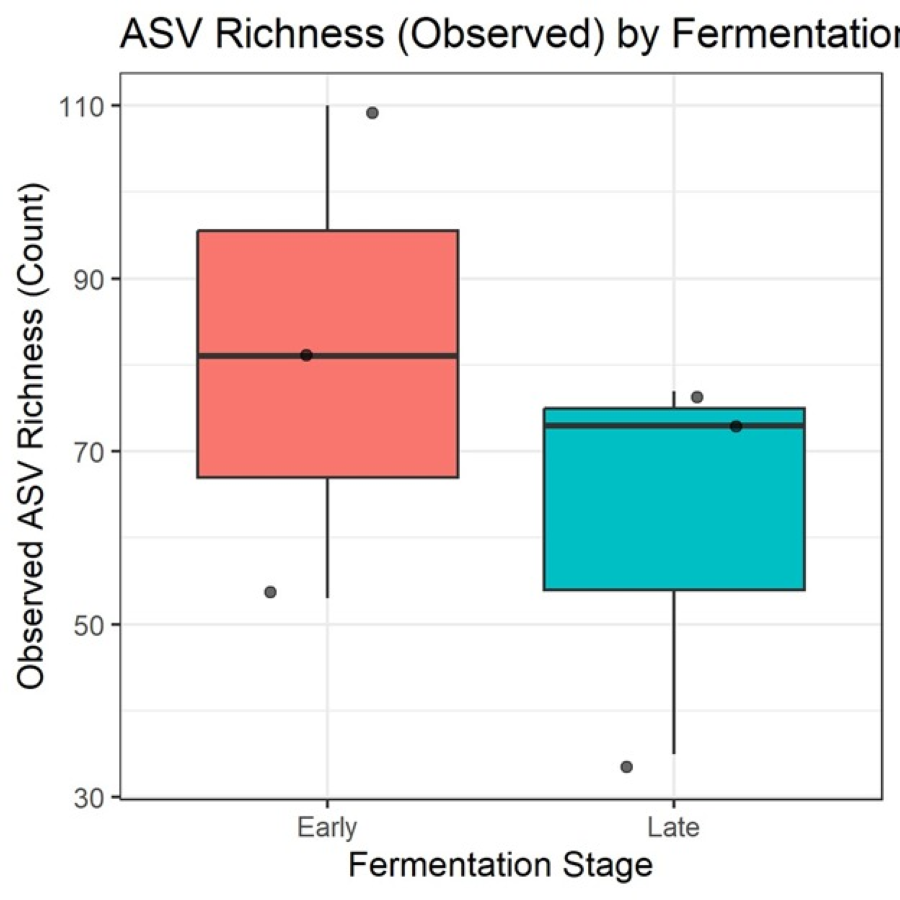

### Supplementary Fig 4

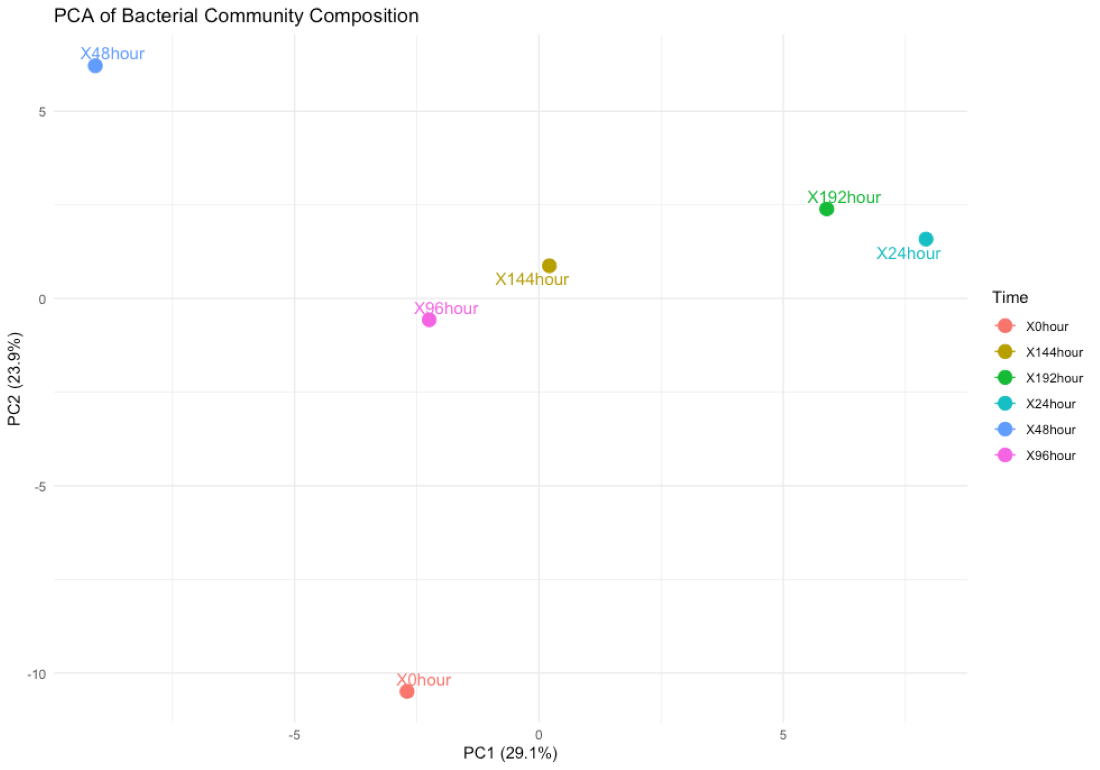

### Supplementary Fig 5

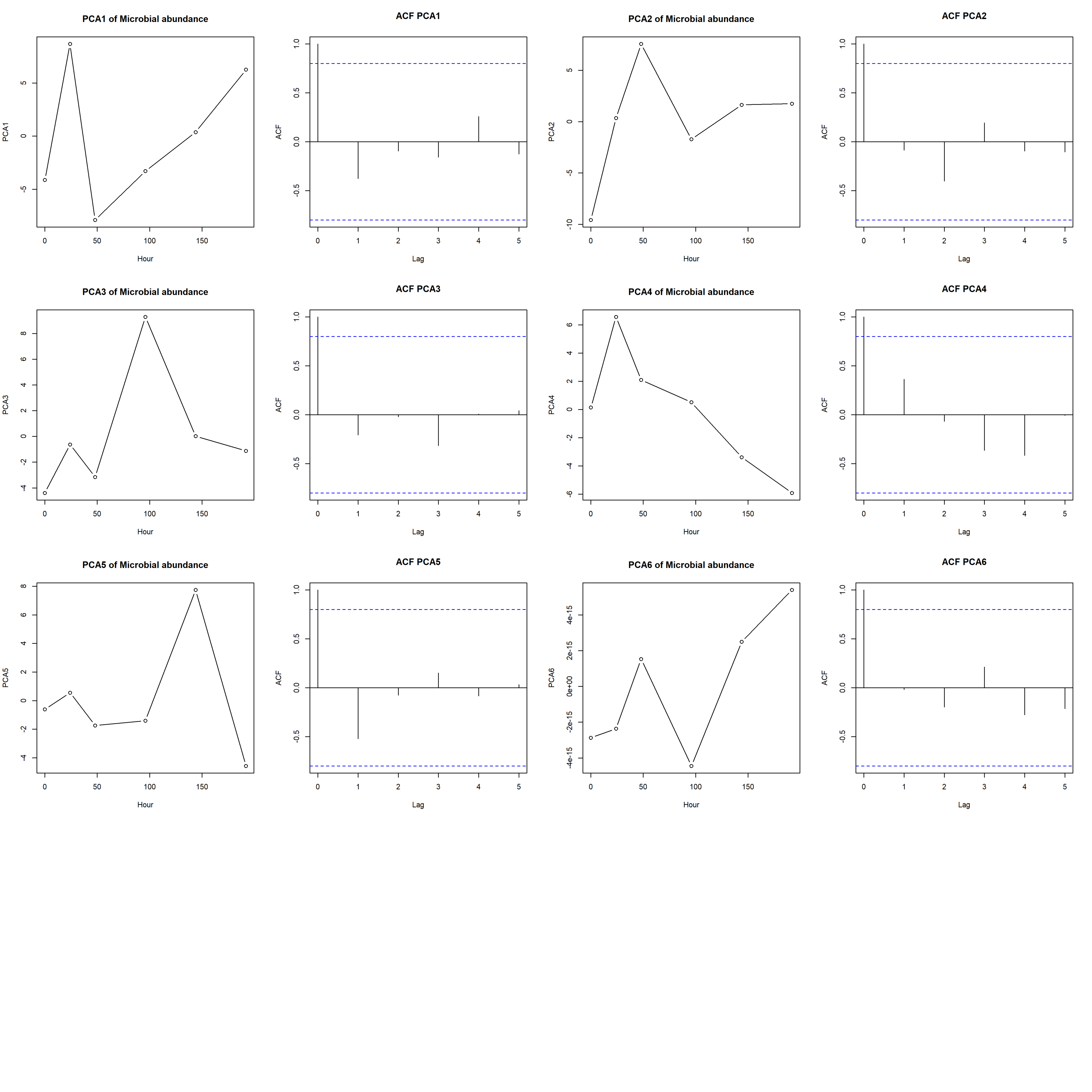

### Supplementary Fig 6

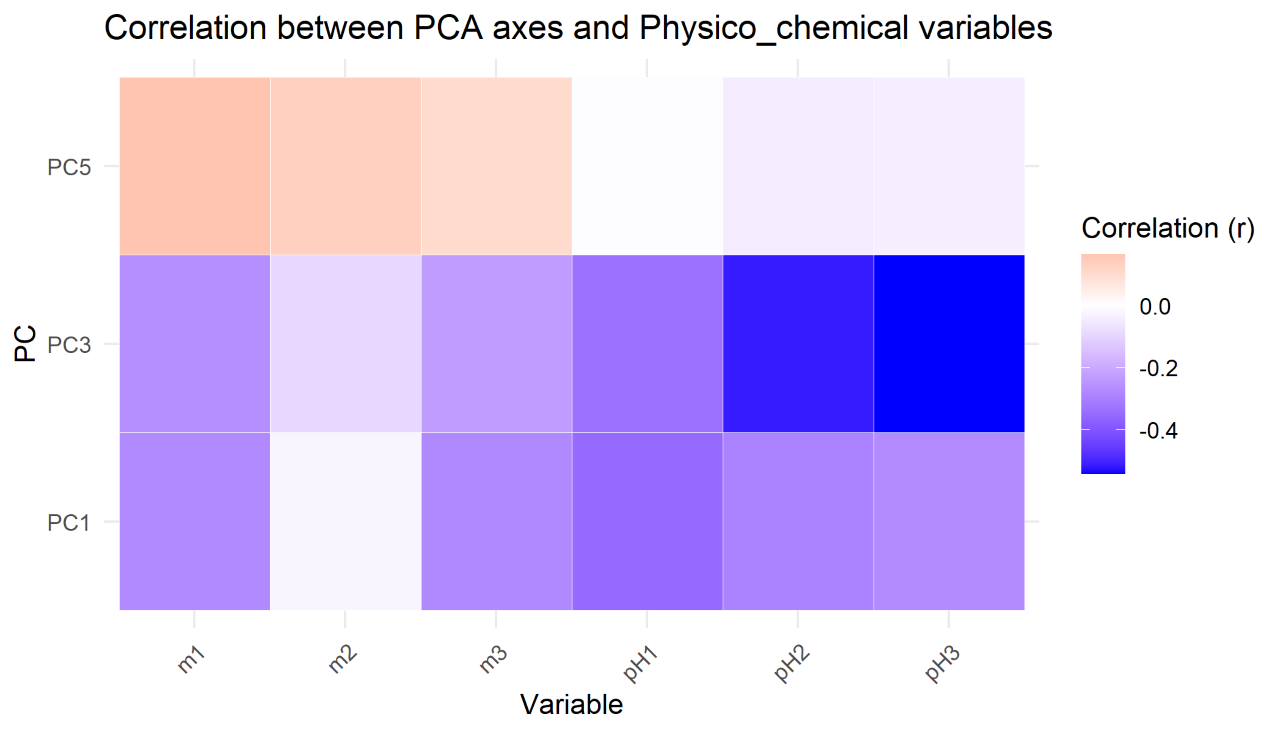

### Supplementary Fig 7

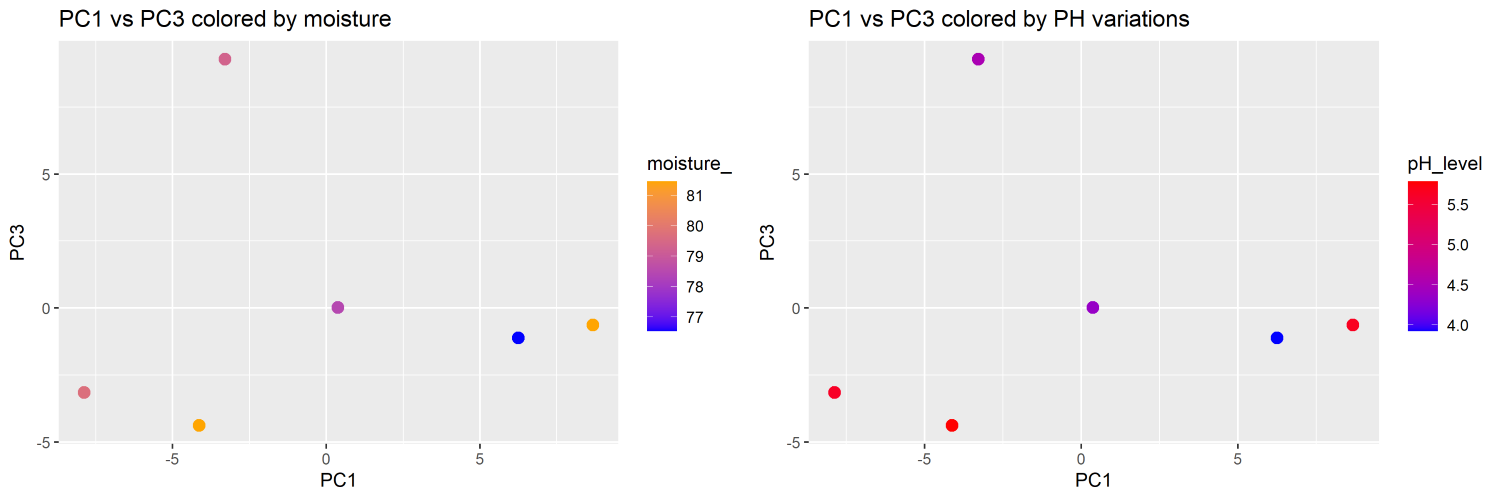
